# Ankyrin-1 gene exhibits allelic heterogeneity in conferring protection against malaria

**DOI:** 10.1101/114959

**Authors:** Hong Ming Huang, Denis C. Bauer, Patrick M. Lelliott, Matthew W. A. Dixon, Leann Tilley, Brendan J. McMorran, Simon J. Foote, Gaetan Burgio

**Affiliations:** Department of Immunology and Infectious Disease, John Curtin School of Medical Research, Australian National University, ACT, Australia; CSIRO, Sydney, NSW, Australia; IFReC Research Building, Osaka University, 3-1 Yamada-oka, Suita, Osaka 565-0871, Japan; Department of Biochemistry and Molecular Biology, Bio21 Institute, Melbourne, Victoria, Australia

**Author notes:** **Corresponding author:** Dr Gaetan Burgio, The John Curtin School of Medical Research, Australian National University, 131 Garran Road, ACT 2601, Australia.

**Keywords:** Ankyrin-1, malaria, erythrocyte cytoskeleton, allelic heterogeneity

## Abstract

Allelic heterogeneity is a common phenomenon where a gene exhibit different phenotype depending on the nature of its genetic mutations. In the context of genes affecting malaria susceptibility, it allowed us to explore and understand the intricate host-parasite interactions during malaria infections. In this study, we described a gene encoding erythrocytic ankyrin-1 (*Ank-1*) which exhibits allelic-dependent heterogeneous phenotypes during malaria infections. We conducted an ENU mutagenesis screen on mice and identified two *Ank-1* mutations, one resulted in an amino acid substitution (MRI95845), and the other a truncated *Ank-1* protein (MRI96570). Both mutations caused hereditary spherocytosis-like phenotypes and confer differing protection against *Plasmodium chabaudi* infections. Upon further examination, the *Ank-1^(MRI96570)^* mutation was found to inhibit intra-erythrocytic parasite maturation, whereas *Ank-1^(MW95845)^* caused increased bystander erythrocyte clearance during infection. This is the first description of allelic heterogeneity in ankyrin-1 from the direct comparison between two *Ank-1* mutations. Despite the lack of direct evidence from population studies, this data further supported the protective roles of ankyrin-1 mutations in conferring malaria protection. This study also emphasized the importance of such phenomenon to achieve a better understanding of host-parasite interactions, which could be the basis of future studies.

## Introduction

Malarial parasites have been co-evolving with humans for thousands of years and have played a major role in shaping the human genome in malaria endemic regions [1,2]. Indeed, many genetic polymorphisms were selected for, as they provide significant host survival advantages during malaria infections [1,3], resulting in high frequencies of protective genetic mutations in malaria endemic regions. The majority of these affect the red blood cells (RBCs), and hence the blood stage of malaria infections [3-5].

Interestingly, these genetic mutations or alleles often exhibit varying degrees of malaria protection even if they affect the same gene, which is influenced by the location and the severity of the mutations [6,7]. This phenomenon, known as “allelic heterogeneity”, is characterized by multiple different phenotypes arising from mutations in a single gene. It has been described for certain genes affecting malaria susceptibility, which is reflected by their geographical distribution within malaria-endemic regions [8]. One of the most prominent examples of this is the G6PD deficiency disorder, which can arise from multiple mutations in the G6PD gene [7,9]. Many studies have explored the effectiveness of each mutation in protecting individuals from malaria, which corresponds to the distribution of each allele across the globe [10-12]. Another example is the β-globin gene, which is well known for its two malaria protective alleles - the HbS and HbC in African populations [13,14]. HbC is restricted to West Africa, whereas HbS is widespread throughout Africa, which is thought to be linked to the effectiveness of each allele at conferring malaria resistance, and their non-malaria-associated morbidity [15,16]. Studies on these alleles would not only allow a better understanding of host-parasite interactions, but also give us insights into the dynamics of population genetics in malaria endemic regions [8].

However, allelic heterogeneity can also complicate the characterization of the malaria protective roles of certain genes, often resulting in conflicting evidence from various studies. One example of such polymorphisms is CD36 deficiency, which was originally thought to be protective against malaria, as evidenced by the positive selection in East Asian and African populations [17-19]. While some studies reported increased malaria protection [20], others reported no significant associations [21] or even increased susceptibility [17,22]. It is possible that these contradictive findings are due to confounding factors associated with allelic heterogeneity in CD36 deficiency [6]. This further emphasizes the importance of taking allelic heterogeneity into consideration better design future studies involving host genetics in malaria, as well as various other infectious diseases.

In terms of malaria susceptibility, however, the allelic heterogeneity of genes affecting RBC cytoskeleton is poorly understood. Many of the resulting genetic disorders are heterogeneous, such as hereditary spherocytosis (HS), which is characterized by the formation of “spherocytic” RBCs that exhibit reduced volume due to disruptions in erythrocyte cytoskeletons. HS is caused by mutations in ankyrin, spectrins, band 3 and protein 4.2, with ankyrin mutations contributing to more than 50% of all HS cases [23-27], where the severity depends greatly on the location and the nature of mutations [28]. However, the prevalence of HS in malaria endemic regions is not well studied; only isolated cases were reported [29-32]. Nevertheless, *in vivo* and *in vitro* studies have repeatedly suggested an association of HS with increased malaria resistance, and several mechanisms have been proposed, although not all of them were consistent [33-36]. Based on these observations, we hypothesized that the inconsistencies in resistance mechanisms might be due to the allelic heterogeneity of genes associated with HS.

To explore this hypothesis, we examined mouse models carrying two novel N-ethyl-N-nitrosourea (ENU)-induced ankyrin mutations. These two mouse lines, *Ank-1^(MRI96570^*^/*+)*^ and *Ank-1(^(MRI95845/MRI95845)^*, displayed hematological and clinical features consistent with HS, and a marked resistance to infection by the murine malarial parasite, *Plasmodium chabaudi*. Analysis of the underlying mechanism of resistance to infection revealed both common and distinct features between the strains. RBCs from both mouse lines were similarly resistant to merozoite invasion. Although, the *Ank-1^(MRI95845/MRI95845)^* erythrocytes were more rapidly cleared from circulation during an infection, an impairment in intra-erythrocytic parasite maturation was observed in the infected *Ank-1^(MRI96570/+)^* erythrocytes. This study highlights the first direct examination of allelic heterogeneity of the *Ank-1* gene in the context of malaria resistance in mouse models.

## Materials and Methods

### Mice and ethics statement

All mice used in this study were housed with 12 hour light-dark cycle under constant temperature at 21 °C, with food and water available *ad libitum*. All procedures were performed according to the National Health and Medical Research Council (NHMRC) Australian code of practice. Experiments were carried out under ethics agreement AEEC A2014/54, which was approved by the animal ethics committees of the Australian National University.

### ENU mutagenesis and dominant phenotype screening

SJL/J male mice were injected intraperitoneally with two doses of 100 mg/kg ENU (Sigma-Aldrich, St Louis, MO) at one week interval. The treated males (G0) were crossed to females from the isogenic background to produce the first generation progeny (G1). The seven-week-old G1 progeny were bled and analyzed on an Advia 120 Automated Hematology Analyzer (Siemens, Berlin, Germany) to identify abnormal red blood cell count. A mouse carrying the MRI96570 or MRI95845 mutation was identified with a RBC “mean corpuscular volume” (MCV) value three standard deviations lower from other G1 progeny. It was crossed with SJL/J mice to produce G2 progeny to test the heritability of the mutations and the dominance mode of inheritance. Mice that exhibited low MCV (<48fL) were selected for whole exome sequencing and genotyping.

### Whole exome seguencing

DNA from two G2 mice per strain carrying the abnormal red blood cell parameters (MCV <48fL) were extracted with Qiagen DNeasy blood and tissue kit (Qiagen, Venlo, Netherlands) for exome sequencing as previously described [37]. Briefly, 10μg of DNA was prepared for exome enrichment with Agilent Sure select kit paired-end genomic library from lllumina (San Diego, CA), followed by high throughput sequencing using a HiSeq 2000 platform. The bioinformatics analysis was conducted according to the variant filtering method previously described by Bauer et al. [38]. Private variants that were shared between the two mutants but not with other SJL/J, C57BL/6 mice, excluding the common variants between other mouse strains were annotated using ANNOVAR [39]. Private non-synonymous exonic and intronic variants within 20 bp from the exon spicing sites were retained as potential candidate ENU mutations.

### PCR and Sanger sequencing

DNA from mutant mice was amplified through PCR with 35 cycles of 30 seconds of 95°C denaturation, 30 seconds of 56-58°C annealing and 72°C elongation for 40 seconds. The primers used in the PCR are described below. The PCR products were examined with agarose gel electrophoresis before being sent to the Australian Genome Research Facility (AGRF) in Melbourne, Australia, for Sanger sequencing. Logarithm of odds (LOD) score was calculated based on the number of mice that segregated with the candidate mutations.

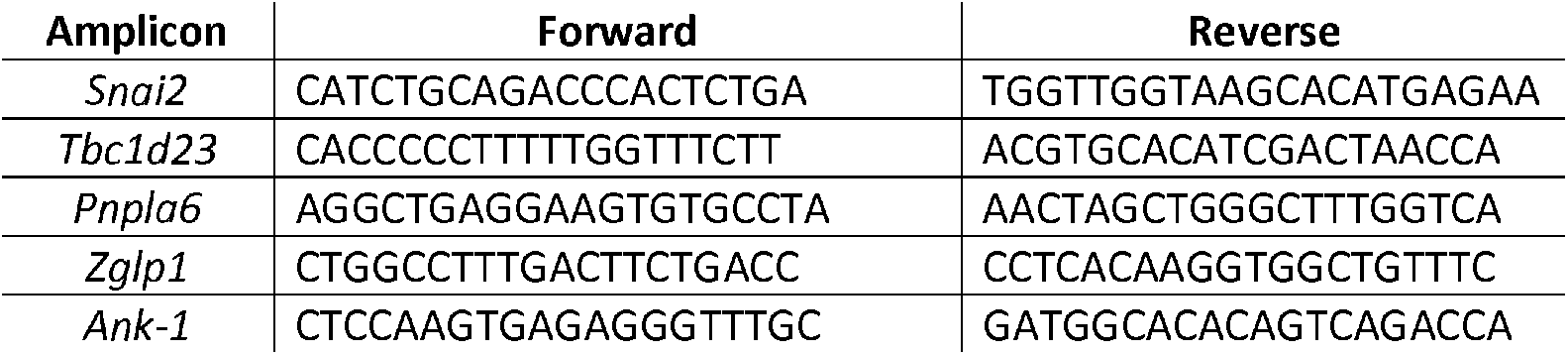
Primers for MRI95845 mutation:

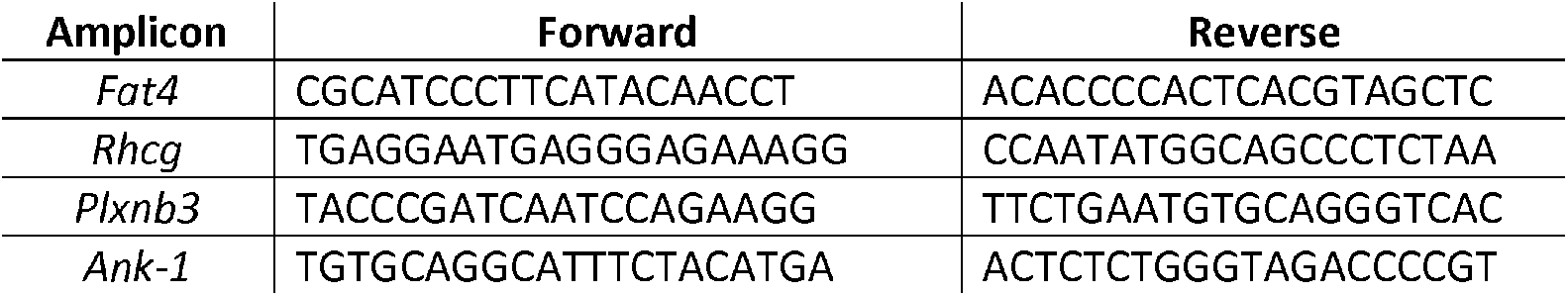
Primers for MRI96570 mutation:

### RBC osmotic fragility analysis

To assess the susceptibility of RBC membrane to osmotic stress, 5μl of mouse whole blood was diluted 100-fold with phosphate buffer (pH 7.4) containing 0 to 10g/L of sodium chloride, and incubated for at least 10 minutes at room temperature. The cells were centrifuged at 800g for 3 minutes, and the supernatant, which contains free hemoglobin, was measured at 540nm to assess the degree of hemolysis. The absorbance values were expressed as percentage of hemolysis, with hemolysis at 0g/L sodium considered as 100% lysis.

### Erythrocyte lifetime assay

Each uninfected mouse was intravenously injected with 1mg of EZ-link^®^ Sulfo-NHS-LC Biotin (Biotin) (Thermo Scientific, Waltham, MA) in mouse tonicity PBS (MT-PBS). 2ul of blood was collected on day 1, 7, 14, 21 and 28 days post-injection. Samples were stained and analyzed using a flow cytometer. The proportion of Biotin-labelled mature RBCs on day 1 was considered as the “starting point” of 100% of labelled cells. For subsequent timepoints, the remaining number of biotin-labelled RBCs were expressed as a percentage of the starting number as the indication of RBC turnover rate.

For infected mice, 1mg of Biotin was injected intravenously as soon as parasitemia was detectable via flow cytometry (approximately 0.05-0.3%). Samples were collected daily and analyzed as above.

### Ektacytometry

10-15ul of uninfected RBCs were first resuspended in 500ul of pre-warmed polyvinylpyrrolidone (PVP) solution at a viscosity of 30 mPa/second at 37 °C until needed. Samples were analyzed according to the manufacturer’s instructions with a RheoScan Ektacytometer (Rheo Meditech, Seoul, South Korea) and the elongation index measured across a range of pressures from 0-20 Pa. Each sample was measured three times to account for technical variabilities. The values were normalized against the wild-type samples.

### In vitro spleen retention assay

The RBC deformability was examined according to the protocol described previous by Deplaine et al. [40] with modifications. Briefly, RBCs from each genotype of mice were stained with 10μg/ml of either hydroxysulfosuccinimide Atto 633 (Atto 633) or hydroxysulfosuccinimide Atto 565 (Atto 565) (Sigma-Aldrich, St Louis, MO), followed by three washes with MTRC (154mM NaCI, 5.6mM KCI, 1mM MgCI_2_, 2.2mM CaCI_2_, 20mM HEPES, 10mM glucose, 4mM EDTA, 0.5% BSA, pH 7.4, filter sterilized). The stained RBCs were mixed in equal proportion and diluted with unstained wild-type RBCs to give approximately 10-20% of the total RBCs being labelled RBCs. The samples were further diluted to 1-2% haematocrit with MTRC, before passing through the filter bed. The prefiltered and post-filtered samples were analysed on BD LSRFortessa (BD Biosciences, Franklin Lakes, NJ) flow cytometer to determine the proportion being retained in the filter bed.

### Scanning electron microscopy (SEM)

SEM was performed as described previously [41]. Mouse blood was fixed overnight in 3% EM-grade glutaraldehyde (Sigma-Aldrich, St Louis, MO) at 4°C immediately upon collection. The samples were washed with MT-PBS 3 times, 10 minute each time. The cells were then adhered to the coverslips with 0.1% polyethylenimine (PEI) for 10 minutes, before washing with MT-PBS. The cells were then dried serially using 30%, 50%, 70%, 80%, 90%, 100%, 100% ethanol, 10 minutes each. The cells were then soaked in 1:1 ethanol: hexamethyldisilazane solution for 10 minutes, followed by 2 washes with 100% hexamethyldisilazane (Sigma-Aldrich, St Louis, MO), each 10 minutes. The coverslips were then air-dried overnight and coated with gold and examined under JEOL JSM-6480LV scanning electron microscope.

### Quantitative PCR and cDNA sequencing

RNA was purified from embryonic livers of E14 embryos using Qiagen RNeasy kit (Qiagen, Venlo, Netherlands), followed by cDNA synthesis using Transcriptor High Fidelity cDNA Synthesis Kit (Roche, Basel, Switzerland), as described previously [41]. Quantitative PCR was performed on ViiA™ 7 Real-Time PCR System (Thermo Scientific, Waltham, MA). The ΔΔC_T_ method [42] was used to determine the cDNA levels of *Ank-1* and the housekeeping gene β-actin and expressed as a fold-change of the mutants to the wild-type. The primers used for *Ank-1* gene spanned exon 2 to 4: *Ank-1*-F: 5’-TAACCAGAACGGGTTGAACG-3’; *Ank-1*-R: 5’-TGTTCCCCTTCTTGGTTGTC-3’; β-Actin-F: 5’-TTCTTTGCAGCTCCTTCGTTGCCG-3’; β-Actin-R: 5’-TGGATGCGTACGTACATGGCTGGG-3’.

### SDS-PAGE, Coomassie staining and Western blot

The analysis of erythrocytic proteins was carried out as described previously [41]. Briefly, RBC ghosts were prepared by lysing mouse RBCs with ice-cold 5mM phosphate buffer (ph7.4) and centrifuging at 20,000g for 20 minutes followed by removal of the supernatant, and repeat until the supernatant became clear. The RBC ghosts or whole blood lysates were denatured in SDS-PAGE loading buffer (0.0625M Tris pH 6.8, 2% SDS, 10% glycerol, 0.1M DTT, 0.01% bromophenol blue) at 95°C for 5 minutes before loading onto a 4-15% Mini-PROTEAN^®^ TGX™ Precast Gels (Bio-Rad, Hercules, CA). The gels were then either stained with Coomassie blue solution (45% v/v methanol, 7% v/v acetic acid, 0.25% w/v Brilliant Blue G) overnight or transferred to a nitrocellulose membrane. The western blot was carried out using these primary antibodies: lug/ml anti-alpha 1 spectrin (clone 17C7), lug/ml anti-beta 1 spectrin (clone 4C3) (Abeam, Cambridge, UK), 0.5ug/ml anti-GAPDH (clone 6C5) (Merck Millipore, Darmstadt, Germany), anti-N-terminal *Ank-1* “p89” at 1:1500 dilution, anti-band 3 at 1:4000 dilution and anti-protein 4.2 at 1:2000 dilution (kind gifts from Connie Birkenmeier, Jackson Laboratory, US). Each primary antibody was detected with the appropriate horseradish peroxidase (HRP)-conjugated secondary antibody at 1:5000 dilution from 1mg/ml stocks. The blots were visualized using ImageQuant LAS 4000 (GE Healthcare Life Sciences, Arlington Heights, IL), and quantified using ImageJ software [43].

### Malaria infection

Malaria infections on mice were performed as described previously [41]. 200μl of thawed *P. chabaudi adami* infected blood was injected into the intraperitoneal cavity of a C57BL/6 donor mouse. When the donor mouse reached 1-10% parasite load (parasitemia), blood was collected through cardiac puncture. The blood was diluted with Krebs’ buffered saline with 0.2% glucose as described previously [44]. Each experimental mouse was infected with 1×10^4^ parasites intraperitoneally. The parasitemia of these mice were monitored either using light microscopy or flow cytometry.

### Terminal deoxynucleotidyl transferase dUTP nick end labelling (TUNEL) staining

The TUNEL assay was carried out as described previously [41] with slight modification. 3μl of infected blood containing 1-10% parasitemia were collected during trophozoite stage and fixed in 1 in 4 diluted BD Cytofix™ Fixation Buffer (BD Biosciences, Franklin Lakes, NJ) for at least day until they were needed. Each sample was then washed twice with MT-PBS, and adhered to a glass slide pre-coated with 0.1% polyethylenimine (PEI) for 10 minutes at room temperature. The excess cells were washed and the glass slide was incubated overnight at room temperature with TUNEL labelling solution (1mM Cobalt Chloride, 25mM Tris-HCI pH 6.6, 200mM sodium cacodylate, 0.25mg/ml BSA, 60uM BrdUTP, 15U Terminal transferase). The slides were washed three times, followed by staining with 50μg/ml of anti-BrdU-FITC antibody (Novus Biologicals, Littleton, CO) in MT-PBT (MT-PBS, 0.5% BSA, 0.05% Triton X-100) for 1 hour. The slides were then washed three times with MT-PBS and mounted with SlowFade^®^ Gold antifade reagent with DAPI (Thermo Scientific, Waltham, MA) and sealed. When the slides were dried, they were examined using Axioplan 2 fluorescence light microscope (Carl Zeiss, Oberkochen, Germany) between 60x to 100x magnification. At least 100 DAPI-positive cells were counted, and each was graded as either positive or negative for TUNEL staining, as an indication of DNA fragmentation.

### In vivo erythrocyte tracking (IVET) assays

The IVET assay was carried out as previously described by Lelliott et al. [45],[46]. Briefly, at least 1.5ml whole blood was collected from mice of each genotype via cardiac puncture, followed by staining with either 10μg/ml of Atto 633 or 125μg/ml of EZ-Link™ Sulfo-NHS-LC-Biotin (Biotin) (Thermo Scientific, Waltham, MA) for 45 minutes at room temperature, followed by washing two times with MT-PBS. The blood was mixed in two different dye combinations to correct for any dye effects. Approximately 1×10^9^ erythrocytes were injected intravenously into infected wild-type mice at 1-5% parasitemia during schizogony stage. Blood samples were collected at various timepoints, from 30 minutes up to 36 hours after injection, and analyzed using flow cytometry. The ratio of infected labelled erythrocytes was determined, as an indication of the relative susceptibility of RBCs to *P. chabaudi* infections. The proportion of labelled blood populations was also tracked over time to determine the clearance of these RBCs from the circulation.

### Flow cytometry analysis of blood samples

For erythrocyte lifetime assays, 2μl of whole blood samples were stained with 2μg/ml streptavidin-PE-Cy7, 1μg/ml anti-CD71-allophycocyanin (APC) (clone R17217), 1μg/ml anti-CD45-APC eFIuor 780 (clone 30-F11) (eBioscience, San Diego, CA), 4μM Hoechst 33342 (Sigma-Aldrich, St Louis, MO) and 12μM JC-1 (Thermo Scientific, Waltham, MA) in MTRC. The samples were washed once with MTRC and further stained with 2μg/ml streptavidin-PE-Cy7 to capture all biotin-labelled cells. Immediately prior to analyzing on flow cytometer, 5μl of 123count eBeads (eBioscience, San Diego, CA) was added to determine the relative levels of anemia.

For both malaria infections and IVET assay, 2μl of whole blood samples were stained with 2μg/ml streptavidin-PE-Cy7 (only for experiments with biotinylated erythrocytes), 1μg/ml anti-CD45-allophycocyanin (APC)-eFluor 780 (clone 30-F11), 1μg/ml anti-CD71 (TFR1)-PerCP-eFluor 710 (clone R17217) (eBioscience, San Diego, CA), 4pM Hoechst 33342 (Sigma-Aldrich, St Louis, MO) and 12μM JC-1 (Thermo Scientific, Waltham, MA) in MTRC. All samples analyzed through flow cytometry were performed on BD LSRFortessa (BD Biosciences, Franklin Lakes, NJ), where 200,000 to 1,000,000 events were collected and visualized on FACSDiva™ and FlowJo software. RBCs were identified by gating on CD71 negative and CD45 negative population, followed by gating on Atto-labelled and Biotin-labelled erythrocytes on appropriate channels (APC for Atto-633, PE for Atto-565 and PE-Cy7 for Biotin). The parasitemia of each labelled erythrocyte population was determined by gating on Hoechst 33342 positive and JC-1 positive population.

### Statistical analysis

The statistical analysis was carried out as described in previous study [41]. Unless otherwise stated, all statistical analysis was carried out using unpaired two-tailed Student’s t-tests without corrections. In particular, the LOD score method coupled with Bonferroni correction was used to determine the causative mutation for MRI96570 and MRI95845, in which the significance threshold and the LOD score for each candidate mutation was calculated, and a LOD score above the threshold was considered statistically significant. The statistical tests for FRAP assay, ektacytometry, RBC lifetime and clearance assays were performed by first calculating the area under curves for each sample, followed by an unpaired Student’s t-test on pooled mice of each genotype. The statistical tests for *in vitro* spleen retention assays were performed using paired Sudent’s t-test, comparing mutant with wild-type RBC population within the same sample. The statistical significance of the malaria survival was tested using the unpaired Log-Rank test on pooled mutant versus wild-type mice. The statistical significance of *Plasmodium* infection was determined via the statmod software package for R (http://bioinf.wehi.edu.au/software/compareCurves) using the ‘compareGrowthCurves’ function with 10,000 permutation, followed by adjustments for multiple testing. The statistical significance for the ratios of IVET assays was determined using the one sample t-test with hypothetical mean of 1.

## Results

### MRI96570 and MRI95845 carry mutations in Ank-1 gene

ENU-treated SJL/J male mice were crossed with wild-type female to produce G1 progeny. The G1 progeny carrying heterozygous point mutations in their genome were then subjected to hematological screening to identify genes affecting RBC properties that might confer malaria protection. Heterozygous G1 mice MRI96570 and MRI95845 were identified from the ENU-dominant screen with mean cellular volume (MCV) three standard deviations below the normal level of the respective parental line: 48.5fl for MRI96570, and 50.6fl for MRI95845, compared to the background of 55.1±1.2fl in SJL/J inbred strain. These mice were crossed with wild-type to produce G2 progeny, where approximately half exhibit low erythrocyte MCV values. Exome sequencing was carried out from two MRI96570 and MRI95845 G2 progeny showing a reduction of MCV to identify the causative genetic mutations. Unique variants shared between these mice were identified and listed in Table S1. A mutation in the ankyrin-1 *(Ank-1)* gene was present in all the affected mice, and co segregated completely with the reduced MCV phenotype for over three generations of crosses. Sanger sequencing revealed a T to A transversion in exon 34 of the *Ank-1* gene for MRI96570 strain, and a T to A transversion in exon 5 of the *Ank-1* gene for MRI95845 strain (Figure S1). They were predicted to cause a nonsense mutation at amino acid position 1398, located in the spectrin-binding domain for MRI96570 mice, and a substitution of tyrosine for asparagine at amino acid residue 149 in the 4^th^ ankyrin repeat for MRI95845 mice (Figure 1A). MRI96570 and MRI95845 will be referred as *Ank-1^(MRI96570)^* and *Ank-1^(MRI95845)^* respectively, for the rest of the report.

**Figure 1.**
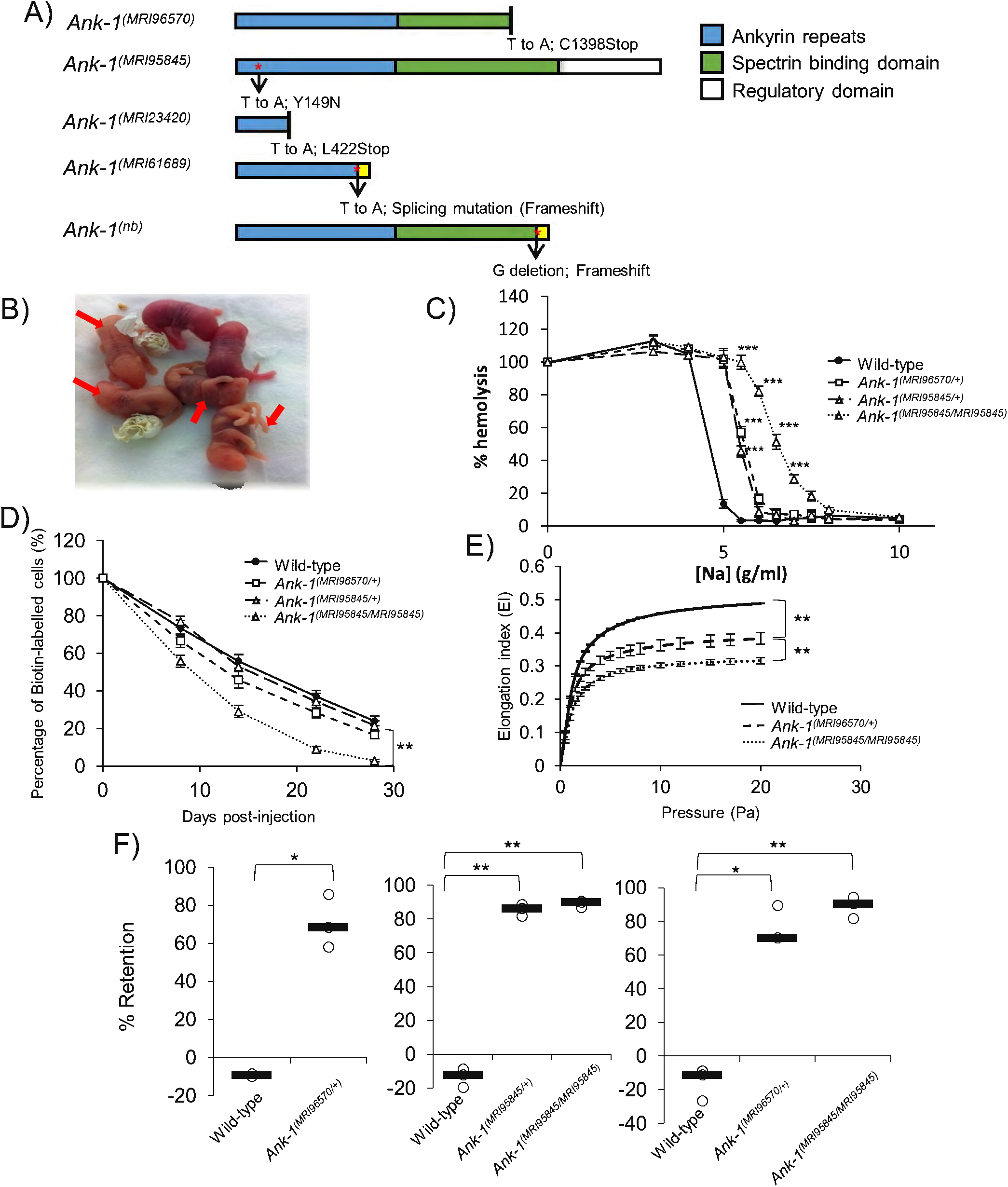
The mutations and phenotypes of *Ank-1^(MRI96570/+)^*, *Ank-1^(MRI95845/+)^* and *Ank-1^(MRI95845/MRI95845)^* mice. (A) The location of ankyrin-1 mutations in *Ank-1^(MRI96570)^* and *Ank-1^(MRI95845/+)^* alleles and the predicted effects on ankyrin-1 protein, compared to the previously described *Ank-1^(MRI23420)^, Ank-1^(MRI61689)^* and *Ank-1^(nb)^*. (B) The *Ank-1^(MRI96570/MRI96570)^* pups (indicated by arrows) showed severe jaundice and died within 1 week after birth. (C) The osmotic fragility of *Ank-1^(MRI96570/+)^, Ank-1^(MRI95845/+)^* and *Ank-1^(MRI95845/MRI95845)^* erythrocytes in hypotonic solution from 0-10g/L sodium (n=5). (D) The RBC half-life of *Ank-1^(MRI96570/+)^, Ank-1^(MRI95845/+)^* and *Ank-1^(MRI95845/MRI95845)^* mice (n=5). (E) The elasticity of *Ank-1^(MRI96570/+)^* and *Ank-1^(MRI95845/MRI95845)^* RBCs under shear pressure as measured by ektacytometer (n=3). (F) The proportion of retained *Ank-1^(MRI96570/+)^, Ank-1^(MRI95845/+)^* and *Ank-1^(MRI95845/MRI95845)^* RBCs when passing through a layer of beads during the *in vitro* spleen retention assay (n=3). * P<0.05, ** P<0.01, *** P<0.001. All error bars indicate standard error of mean (SEM).

### Both *Ank-1^(MRI96570)^* and *Ank-1^(MRI95845)^* exhibit HS-like phenotypes

Since ankyrin mutations are usually associated with HS, we examined both *Ank-1^(MRI96570)^* and *Ank-1^(MRI95845)^* mice in terms of their HS-like phenotypes. When two *Ank-1^(MRI96570/+^)* G2 progeny were intercrossed, *Ank-1^(MRI96570/MRI96570)^* mice were born with severe jaundice and died within several days of birth (Figure 1B), suggesting homozygosity for *Ank-1^(MRI96570)^* mutation caused lethal anemia. On the other hand, *Ank-1^(MRI95845/MRI95845)^* mice appeared healthy with a normal lifespan. Hematological analysis of these mice revealed a significant reduction in MCV and mean corpuscular hemoglobin (MCH), and increased red cell distribution width (RDW) (Table S2), indicating microcytosis and anisocytosis, which are the hallmarks for HS. When the RBCs were subjected to osmotic stress, RBCs from *Ank-1^(MRI96570^*^/*+)*^, *Ank-1^(MRI95845^*^/*+)*^ and *Ank-1^(MRI95845/MRI95845)^* mice exhibit significantly increased osmotic fragility compared to wild-type RBCs (Figure 1C). In particular, sodium chloride concentration required to achieve 50% hemolysis is significantly higher (P<0.001) for *Ank-1^(MRI96570/+)^* RBCs (5.6 g g/L or 104mM) and *Ank-1^(MRI95845/+)^* RBCs (5.4 g/L or 100mM), compared to wild-type RBCs (4.6 g/L or 84mM). The *Ank-1^(MRI95845/MRI95845)^* RBCs showed further susceptibility to osmotic stress, with 50% hemolysis at approximately 6.5 g/L (121mM) sodium chloride concentration.

We predicted that the mutant RBCs have shorter half-life, which is also one of the symptoms of HS. Therefore, RBC half-life was determined by biotinylating mouse RBCs *in situ* and tracking the proportion of biotinylated RBCs in circulation over time. As shown in Figure 1D, erythrocytes from *Ank-1^(MRI95845/MRI95845)^* have a Significantly shorter half-life of approximately 9.5 days as opposed to the 16 days of wild-type erythrocytes (P=0.008), but no significant difference was observed for erythrocytes from heterozygous mice (P=0.09 for *Ank-1^(MRI96570^*^/*+)*^ mice and P=0.08 for *Ank-1^(MRI95845^*^/*+)*^ mice). The morphology of these RBCs were examined under light and scanning electron microscopy (Figure S2). *Ank-1^(MRI96570^*^/*+)*^ and *Ank-1^(MRI95845^*^/*+)*^ mice exhibited slight reduction in RBC size, while *Ank-1^(MRI95845/MRI95845)^* mice had smaller acanthocytic RBCs and displayed anisocytosis. On the other hand, blood smears obtained from jaundiced *Ank-1^(MRI96570/MRI96570)^* pups showed reticulocytosis, fragmented RBCs and severe anisocytosis.

Another feature of HS is reduced RBC deformability, which was examined using two different analytical techniques: ektacytometry and an *in vitro* spleen retention assay. Ektacytometry measures the flexibility of RBCs when subjected to shear pressure, and expresses as an elongation index, which indicates the deformability of RBCs. The *Ank-1^(MRI96570/+)^* RBCs showed reduced elongation index compared to wild-type, with *Ank-1^(MRI95845/MRI95845)^* RBCs showing further reduction in elongation index, indicating significant reduction in RBC deformability (Figure 1E). In addition, the *in vitro* “spleen mimic” retention assay was performed by passing the erythrocytes through layer of microbeads of varying sizes, modelling *in vivo* splenic filtration. RBC deformability was assessed by the ability of RBCs to pass through the bead layer. Figure 1F showed three independent measurements of RBC deformability using the splenic retention assay, comparing wild-type, *Ank-1^(MRI96570/+)^*, *Ank-1^(MRI95845/+)^* and *Ank-1^(MRI95845/MRI95845)^* RBCs. approximately 70% increased retention for *Ank-1^(MRI96570/+)^* RBCs was observed compared to wild-type, whereas erythrocytes of *Ank-1^(MRI95845/+)^* and *Ank-1^(MRI95845/MRI95845)^* mice showed 86% and 90% increased RBC retention compared to wild-type, respectively. However, no significant difference was observed between *Ank-1^(MRI96570/+)^* and *Ank-1^(MRI95845/MRI95845)^* erythrocytes.

The expression levels of ankyrin and other RBC membrane proteins were also examined (Figure S3). Significant reduction of *Ank-1* mRNA levels was observed in *Ank-1^(MRI96570/+)^*, *Ank-1^(MRI95845/+)^*, *Ank-1^(MRI96570/MRI96570)^* and *Ank-1^(MRI95845/MRI95845)^* embryonic livers (Figure S3a). However, Coomassie staining and Western blotting of the RBC membrane fractions did not show a significant difference in ANK-1 levels between wild-type, *Ank-1^(MRI96570/+)^* and *Ank-1^(MRI95845/MRI95845)^* erythrocytes (P=0.3) (Figure S3b-d), using an anti-ANK-1 antibody (p89) targeting specifically to the N-terminal region of ANK-1 protein [36]. The predicted truncated ANK-1^*(MRI96570/+)*^ form (160kDa) was also not evidenced, suggesting degradation of truncated protein. The levels of other cytoskeletal proteins were also examined to account for possible disruptions to interactions with binding partners of ankyrin-1. However, no difference was observed for band 3, α- and β-spectrin, whereas with a significantly lower protein 4.2 level was observed only in *Ank-1^(MRI95845/MRI95845)^* erythrocytes (Figure S3d).

### *Ank-1^(MRI96570)^* and *Ank-1^(MRI95845)^* confer protection against *P. chabaudi* infection

We proposed that mice carrying these mutations have reduced susceptibility to malaria infection, which we examined by injecting with a lethal dose of *P. chabaudi*, and recording the percentage of parasitized RBCs (parasitemia). As shown in Figure 2A, *Ank-1^(MRI96570^*^/*+)*^ and *Ank-1^(MRI95845^*^/*+)*^ mice showed significant reduction in peak parasitemia of approximately 15-20%, while *Ank-1^(MRI95845/MRI95845)^* mice showed approximately 30% reduction in peak parasitemia compared to wild-type. *Ank-1^(MRI95845/MRI95845)^* mice also showed a two-day delay in parasitemia, peaking on day 12 post-infection rather than day 10 as with wild-type. *Ank-1^(MRI95845/MRI95845)^* mice also exhibited significantly higher survival rate compared to wild-type during *P. chabaudi* infection, but no significant difference was observed for *Ank-1^(MRI96570^*^/*+)*^ and *Ank-1^(MRI95845^*^/*+)*^ mice compared to wild-type (Figure 2B). Overall, these results suggested that both *Ank-1^(MRI96570^*^/*+)*^ and *Ank-1^(MRI95845^*^/*+)*^ mice showed moderate resistance, whereas *Ank-1^(MRI95845/MRI95845)^* mice exhibited significant resistance towards *P. chabaudi* infection relative to wild-type mice.

**Figure 2.**
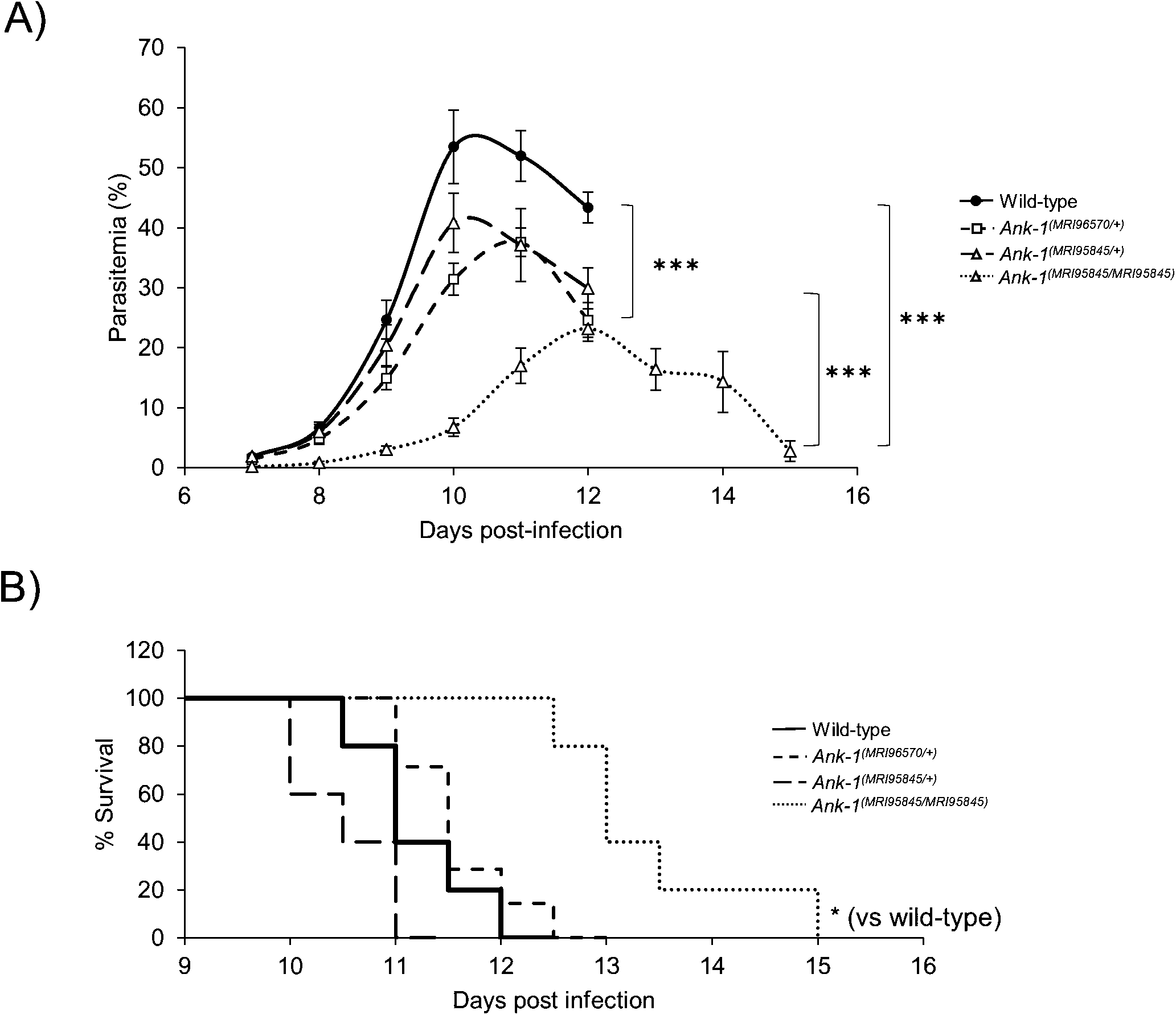
The parasitemia and survival of *Ank-1^(MRI96570/+)^, Ank-1^lMW95845/+)^* and *Ank-1^(MRI95845/MRI95845)^* mice during *P. chabaudi* infection. (A) The parasite load and (B) survival rate of *Ank-1^(MRI96570^*^/*+)*^, *Ank-1^(MRI95845)^* and *Ank-1^(MRI95845/MRI95845)^* mice from two independent experiments, starting from day 7 to day 15 post-infection when challenged with 1x10^4^ parasite intraperitoneally as determined by light microscopy. (n=9-13). * P<0.05, *** P<0.001. Error bars indicate SEM.

From these results, we further investigated and compared the possible mechanisms of resistance mediated by *Ank-1^(MRI96570)^* and *Ank-1^(MRI95845)^* mutations. We examined three important determinants of parasite growth and survival within the host. Firstly, we studied the ability of parasite to survive within these erythrocytes, since ankyrin-1 mutations have previously been implicated to impair parasite intra-erythrocytic maturation [36]. Secondly, the erythrocyte invasion was assessed since the mutations disrupt erythrocyte cytoskeletal structure, which is important for facilitating efficient erythrocyte invasion [47]. Thirdly, the mutations might result in an improved detection of parasitized RBCs, thus enhancing their removal from circulation during malaria infection. Since *Ank-1^(MRI96570/+)^* and *Ank-2^(MRI95845/MRI95845)^* mice exhibited differences in malaria resistance, we hypothesized that they mediate malaria resistance through different pathways.

### *Ank-1^(MRI96570/+)^* and *Ank-1^(MRI95845/MRI95845)^* erythrocytes are resistant to merozoite invasion

First, the ability of parasites to invade erythrocytes was assessed via an *in vivo* erythrocyte tracking (IVET) assay. Labelled RBCs from either wild-type, *Ank-1^(MRI96570/+)^* or *Ank-1^(MRI95845/MRI95845)^* mice were injected into infected wild-type mice of 1-10% parasitemia during late schizogony stage and the parasitemia of each genotype was monitored over 36-40 hours to indicate relative invasion rates. The initial invasion period was expected at 30 minutes to 3 hours post-injection, and the results were expressed as a ratio of parasitized RBCs of either, *Ank-1^(MRI96570/+)^* to wild-type (Figure 3A), *Ank-1^(MRI95845/MRI95845)^* to wild-type (Figure 3B), or *Ank-1^(MRI96570/+)^* to *Ank-1^(MRI95845/MRI95845)^*(Figure 3C) From Figure 3A and 3B, *Ank-1^(MRI96570/+)^* and *Ank-1^(MRI95845/MRI95845)^* erythrocytes were less parasitized compared to wild-type (0.6-0.7 for *Ank-1^(MRI96570/+)^* to wild-type, P<0.001; and 0.55-0.8 for *Ank-1^(MRI95845/MRI95845)^* to wildtype, P<0.001) from 3 hours up to 36 hours post-injection, indicating both *Ank-1^(MRI96570/+)^* and *Ank-1^(MRI95845/MRI95845)^* erythrocytes were more resistant to parasite invasion than wild-type. However, no significant differences in parasitemia ratio were observed at the 30 minute timepoint. Furthermore, when the invasion rate of both *Ank-1^(MRI96570/+)^* and *Ank-1^(MRI95845/MRI95845)^* erythrocytes were compared in infected wild-type mice (Figure 3C), no significant difference in parasitemia ratio was observed, suggesting a similar invasion rate between the two mutant erythrocytes.

**Figure 3.**
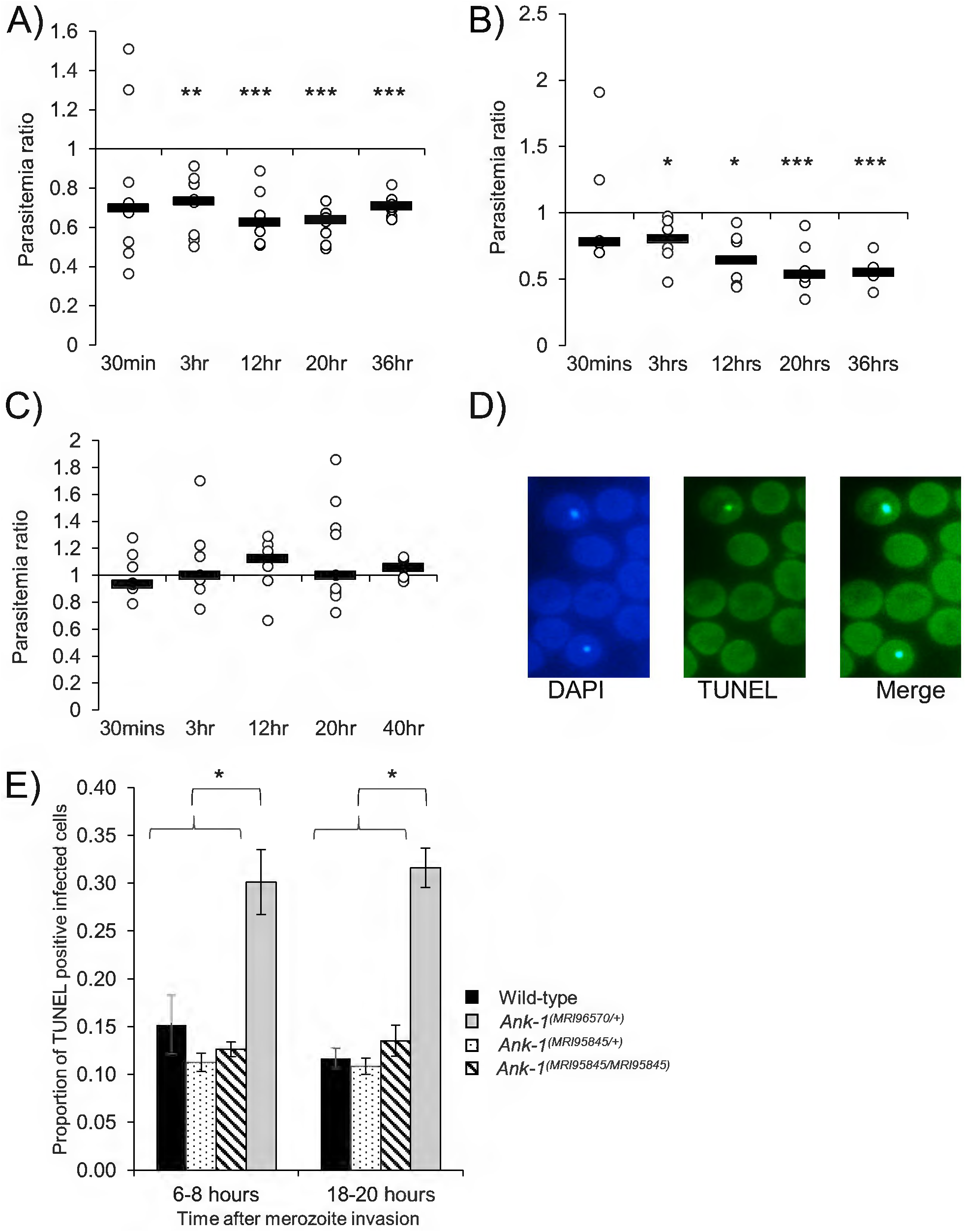
The parasite invasion and intra-erythrocytic growth as indicated via IVET and TUNEL assay. The relative invasion efficiency into *Ank-1^(MRI96570/+)^* and *Ank-1^(MRI95845/MRI95845)^* erythrocytes was examined through IVET assay, where parasitemia ratio was calculated from parasite load of either (A) *Ank-1^(MRI96570/+)^* to wild-type, (B) *Ank-1^(MRI95845/MRI95845)^* to wild-type, or (C) *Ank-1^(MRI96570/+)^* to *Ank-1^(MRI95845/MRI95845)^* erythrocytes (n=5-7 per group). (D) The parasite growth inhibition was determined via TUNEL assay on infected RBCs (DAPI-positive) as an indicator of apoptotic and necrotic parasites. (E) The proportion of TUNEL-positive infected RBCs was counted for *Ank-1^(MRI96570/+)^, Ank-1^(MRI95845/+)^* and *Ank-1^(MRI95845/MRI95845)^* mice at 1.5% parasitemia at ring stage (6-8 hours) and late trophozoite (1820 hours) stage (n=4). *P<0.05, **P<0.01, ***P<0.001. Error bars indicates SEM.

### *Ank-1^(MRI96570/+)^* erythrocytes impair parasite maturation

Second, the parasite intra-erythrocytic maturation was determined through a TUNEL assay, which allows the detection of fragmented DNA in RBCs, as an indication of dying parasites (Figure 3D) [48]. Samples were collected from infected mice at 1-10% parasitemia at both young ring stage and late trophozoite stage, and the proportion of TUNEL-positive infected RBCs were measured. As seen from Figure 3E, more TUNEL-positive parasites were observed within *Ank-1^(MRI96570^*^/*+)*^ erythrocytes, in both ring (30.1±3.4% compared to 15.2±3.1% of wild-type) and trophozoite stage (30.8±3.8% compared to 11.7±1.0% of wild-type), whereas no differences were observed for *Ank-1^(MRI95845^*^/*+)*^ and *Ank-1^(MRI95845/MRI95845)^* erythrocytes. This result suggested that the growth of parasites within *Ank-1^(MRI96570^*^/*+)*^ erythrocytes was impaired, but was normal in *Ank-1^(MRI95845^*^/*+)*^ and *Ank-1(MRI95845/MRI95845)* erythrocytes. This also indicate that *Ank-1^(MRI96570)^* disrupts parasite maturation, whereas *Ank-1^(MRI95845)^* seems to support normal parasite growth.

### *Ank-1^(MRI95845/MRI95845)^* erythrocytes are more likely to get cleared during malaria infections, partially via splenic filtration

The proportions of labelled erythrocytes were also monitored during the IVET assays to compare the relative loss of the two labelled RBC populations as an indicator of RBC clearance during malaria infection. No significant reduction in *Ank-1^(MRI96570^*^/*+)*^ erythrocyte numbers was observed during IVET assay compared to wild-type (Figure 4A). In contrast, the number of labelled *Ank-1^(MRI95845/MRI95845)^* erythrocytes decreased significantly compared to wild-type and *Ank-1^(MRI96570^*^/*+)*^ erythrocytes (Figure 4B and C), with approximately 20% and 50% reduction, respectively. However, the parasitemia measurements during the IVET assays were approximately 2% to 16-30% (Figure S4a-b), which did not correlate with the reduction of labelled *Ank-1^(MRI95845/MRI95845)^* erythrocytes. This suggested an increased bystander clearance rather than clearance of infected *Ank-1^(MRI95845/MRI95845)^* RBCs. To further verify this observation, the infected mice from each genotype were injected with biotin and the biotinylated RBC half-life was examined. As shown in Figure 4D, the *Ank-1^(MRI96570^*^/*+)*^ mice exhibited no significant reduction in RBC numbers, whereas *Ank-1^(MRI95845/MRI95845)^* mice were found to have significantly shorter half-life of approximately 6 days, which did not correlate with the parasitemia curve (Figure S4c). This observation of shorter RBC half-life in infected *Ank-1^(MRI95845/MRI95845)^* mice is consistent with the increased *Ank-1^(MRI95845/MRI95845)^* erythrocyte clearance as shown in IVET assays.

**Figure 4.**
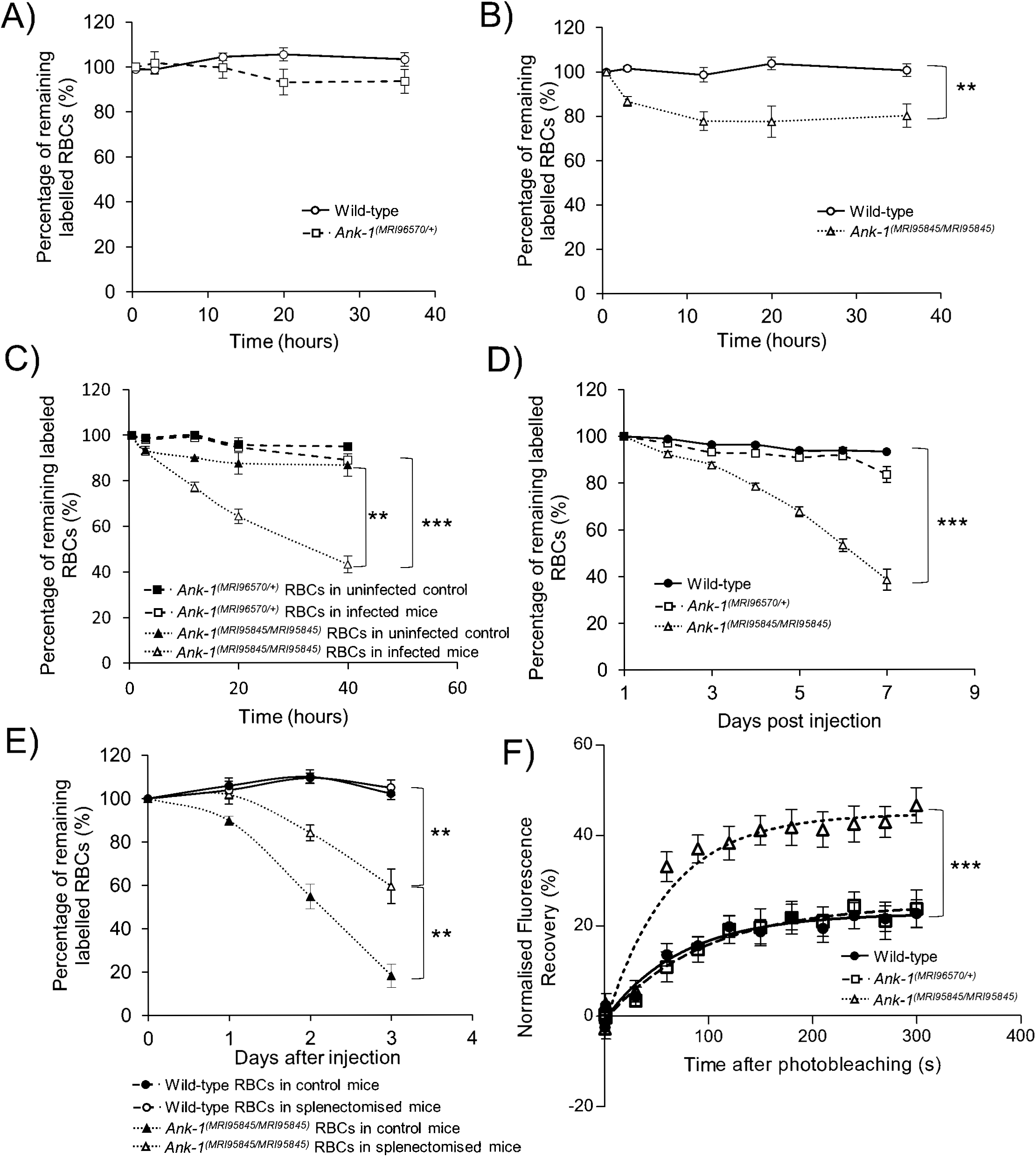
The band 3 mobility and clearance of wild-type, *Ank-1^(MRI96570/+)^* and *Ank-1^(MRI95845/MRI95845)^* erythrocytes. The remaining percentage of labelled RBCs was monitored during the course of IVET assays, comparing between (A) wild-type and *Ank-1^(MRI96570/+)^* erythrocytes, (B) wild-type and *Ank-1^(MRI95845/MRI95845)^* erythrocytes, and (C) *Ank-1^(MRI570/+)^* and *Ank-1^(MRI95845/MRI95845)^* erythrocytes (n=5-7). (D) The half-life of wild-type, *Ank-1^(MRI96570/+)^ Ank-1^(MRI95845/MRI95845)^* erythrocytes during malaria infection as determined by biotinylation of RBCs when parasites were detectable (n=6-7). (E) The clearance of wild-type and *Ank-2^(MRI95845/MRI95845)^* erythrocytes in splenectomised and non-splenectomised mice infected with *P. chabaudi* over 3 days starting from 1% parasitemia (n=6). (F) The band 3 mobility on RBC membrane was measured using Fluorescence recovery after Photobleaching (FRAP), showing the recovery rate of fluorescence as a result of Band 3 migration to the bleach spot (n=9-21). ** P<0.01, *** P<0.001. Error bars indicate SEM.

We proposed that the spleen played a major role in mediating this bystander clearance. Therefore, we infected splenectomized mice with *P. chabaudi* and infused with labelled wild-type and *Ank-1^(MRI95845/MRI95845)^* erythrocytes, the proportions of which were monitored over time. As shown in Figure 4E, *Ank-1^(MRI95845/MRI95845)^* erythrocyte numbers are approximately two-fold higher (P<0.01) in splenectomized mice compared to non-splenectomized mice. This suggests that the spleen is a major contributor towards *Ank-1^(MRI95845/MRI95845)^* erythrocyte clearance, although the clearance was not completely abrogated in the absence of the spleen.

### Increased Band 3 mobility in *Ank-1^(MRI95845/MRI95845)^* erythrocytes as a likely mechanism for increased clearance

Nevertheless, we hypothesized that this increase in RBC clearance is likely due to changes to the cytoskeletal structure of *Ank-1^(MRI95845/MRI95845)^* RBCs. In order to examine our hypothesis, we investigated the band 3 mobility within the RBC membrane as an indicator of the integrity of vertical linkages in the RBC cytoskeleton [49,50]. We fluorescently labeled erythrocytic band 3 with eosin-5’-maleimide and performed Fluorescence Recovery after Photobleaching (FRAP) on erythrocytes, which involves photobleaching a spot on the RBC surface with a pulse from a high-powered laser followed by a fluorescence recovery period where the intensity was recorded over 5 minutes as a marker of mobility. *Ank-1^(MRI95845/MRI95845)^* RBCs were found to have significantly higher fluorescence recovery compared to wild-type and *Ank-1^(MRI96570^*^/*+)*^ RBCs (Figure 4F), which suggests a higher band 3 mobility in *Ank-1^(MRI95845/MRI95845)^* erythrocytes, possibly due to an increased amount of band 3 that was not associated with the RBC cytoskeleton as the result of disrupted ankyrin binding to band 3 on RBC surface.

## Discussion

### *Ank-1* gene displayed allele-dependent heterogeneous phenotypes during malaria infections

Similar to HS in human populations, ankyrin mutations in mice also exhibit differences in clinical symptoms depending on the mutations. As shown in this study, homozygosity for MRI96570 mutation is lethal, while MRI95845 homozygotes appeared healthy; whereas both *Ank-1^(MRI96570^*^/*+)*^ mice and *Ank-1^(MRI95845^*^/*+)*^ mice exhibited HS-phenotypes with similar severity. While both mutations also conferred malaria protection and appeared to impair parasite invasion, they also showed some remarkable differences in mediating this resistance. Parasites in *Ank-1^(MRI96570^*^/*+)*^ erythrocytes were more likely to be TUNEL-positive, indicating impaired intra-erythrocytic maturation, whereas *Ank-1^(MRI95845/MRI95845)^* erythrocytes were more likely to be removed from circulation and possibly had an increased turnover rate.

These findings were not exclusive to the two *Ank-1* mice described in this study. In fact, previous studies on other *Ank-1* mice also exhibit similar mechanisms of resistance. Notably, similar to *1^(MRl96570^*^/*+)*^, *Ank-1^(MRI23420^*^/*+)*^ [36] and *Ank-1^(nb/nb)^* mice [33] were both reported to affect the parasite survival within the defective RBCs. These mutations resulted in truncated protein, therefore, it is possible that the loss of C-terminal ANK-1 protein might be important for growth. However, *Ank-1^(MRI61689^*^/*+)*^ mice, which were also predicted to give rise to truncated protein, were found to exhibit increased RBC bystander clearance but no intraerythrocytic growth impairment [41], similar to the *Ank-1^(MRI95845/MRI95845)^* mice in this study.

Taking these findings together, it would seem that although these mutations resides in different exons of ankyrin-1 gene, they all resulted in HS-like phenotype. However, they exert different effects on the parasite survival depending on the nature of the mutation. In particular, nonsense mutations (*Ank-1^(nb)^, Ank-1^(MRI23420)^, Ank-1^(MRI96570)^*), with the exception of *Ank-1^(MRI61689)^*, impair parasite growth. On the other hand, the only described missense mutation, *Ank-1^(MRI95845)^*, increases the bystander RBC clearance. It is interesting to note that although *Ank-1^(MRI61689)^* mutation was predicted to produce a truncated protein, a full length alternative spliced transcript with a skipped exon was also found [41], possibly indicating that *Ank-1^(MRI61689)^* is unlikely to be a null mutation, consequently exhibiting different mechanisms of malaria resistance compared to other nonsense mutations. Therefore, it is proposed that the presence of a truncated form of ANK-1 protein might be important factor to determine the detrimental effects on malaria parasites.

However, without detailed examination into RBC cytoskeletal structure, it is challenging to speculate the exact mechanisms of malaria resistance of these mutations. Nevertheless, this is the first direct report of such allelic heterogeneity described in *in vivo* malaria mouse models, highlighting the complexity underlying the genetic resistance to malaria, which is likely to correlate with human populations.

### Allelic heterogeneity *Ank-1* and its association with malaria

Due to lack of large scale studies on the HS prevalence in malaria endemic regions, ankyrin-1 mutations have not been associated with malaria protection. Although HS prevalence is more well-characterized in non-malarial regions such as Northern European and Japanese populations, with a prevalence of about 1 in 2000 [51-53], one study proposed an increased HS incidence in Algeria of about 1 in 1000 [54], raising the possibility of positive selection of HS by malarial parasites. However, as the result of extreme allelic heterogeneity of HS-causing genes, many alleles do not reach sufficient frequencies [55] or achieve consistent symptoms [56] to be easily associated with malaria protection. In addition, technical difficulties [29], confounding factors from large genetic variation in African populations [57], as well as poor diagnostics and health systems [57], pose significant challenges for dissecting the connection between HS and malaria. Furthermore, the varying allele frequency in African populations might introduce epistasis effects, possibly masking the genotype-phenotype associated typically observed in other populations [58]. With development of more advanced technologies and better characterization of the genetic structure of African populations, further studies into the association of HS and malaria could potentially yield beneficial insights into the co-evolutionary relationships between humans and *Plasmodium*.

Nonetheless, previous *in vivo* studies have indicated that *Ank-1* mutations affect merozoite invasion and maturation [33,36], both of which were also demonstrated in this study. However, this study also describes for the first time, the direct *in vivo* observation of different mutations in the *Ank-1* gene mediating two distinct, independent mechanisms of malaria resistance, where one impairs parasite maturation and the other increases RBC clearance. Ankyrin is one of the key proteins involved in RBC remodeling by parasites [59-61], and maintaining the native structure of the RBC cytoskeleton [28,62]. It is possible that this allelic heterogeneity is due to the effect each mutation has on the integrity of the RBC cytoskeletal structure, as evidenced by the significantly increased band 3 mobility caused by *Ank-1^(MRI95845)^*, but not *Ank-1^(MRI96570)^* mutation (Figure 4F). This suggests that mutations at different locations of the ankyrin-1 protein might have different effects on the RBCs, consequently impacting the ability of parasites to invade and grow, which could be the basis for further studies, while also taking into potential confounding factors due to differences in genetic background.

### Similarities of allelic heterogeneity in *Ank-1* and other malaria susceptibility genes

As evidenced from this study, the protective effect of the *Ank-1* gene against malaria is dependent on the nature and the location of mutations within the gene. Similarly, this allelic heterogeneity is also observed in other malaria susceptibility genes in human populations. For instance, although many G6PD deficiency-causing alleles have been implicated with malaria protection [63,64], the protective effects are often debated, with many studies reporting increased, or no protection, for individuals with certain alleles of G6PD deficiency [65-70]. This is thought to be due to the phenotypic complexity of G6PD deficiency associated with malaria susceptibility [7]. Indeed, various G6PD alleles have been shown to cause a reduction of G6PD activity with differing severity, and were proposed to correlate with the malaria protection they conferred [65]. More recently, Clarke and colleagues proposed reduced G6PD activity is associated with lower risk of cerebral malaria, but in exchange for higher risk of malarial anemia [12], suggesting a delicate balance underlying the high frequency of G6PD polymorphism of populations in malaria endemic region. Similarly, *Ank-1* mutations described in this study, as well as other previous mouse studies [33,35,36], exhibit variability in malaria resistance, most likely as the result of allelic heterogeneity.

The heterogeneity in malaria resistance mechanisms of the *Ank-1* gene as observed in this study is comparable to the two prevalent alleles of the [β-globin gene - the HbS and HbC, which result from amino acid substitution at position 6 from glutamate to either valine, or lysine, respectively. These two mutations exhibit some similarities in mediating malaria resistance, including impaired parasite growth [71,72], reduced cytoadherence [73-75] and increased erythrocyte clearance [76]. However, HbS erythrocytes were found to be more resistant to all forms of malaria, whereas HbC erythrocytes appeared to be protective against cerebral malaria [15]. This difference in malaria protection was proposed to correlate with distribution of HbS and HbC in Africa [16], further emphasizing the importance of allelic heterogeneity in understanding host-parasite interactions.

In conclusion, we have reported a novel observation where the *Ank-1* gene exhibits phenotypic heterogeneity in malaria resistance mechanisms, either by impairing intra-erythrocytic parasite growth, or promoting RBC clearance. This study also highlighted that the allelic heterogeneity in relation to malaria resistance is not exclusive to G6PD deficiency, and it could also be much more common than we expected. Further studies should extend the understanding of the effects of various *Ank-1* mutations on the development of intra- erythrocytic parasites, as well as the association of HS with malaria in human populations. This could provide new insights into the intricate relationships between RBC cytoskeletal structure and parasite interactions.

## Authors’ Contributions

H.M.H., D.C.B., P.M.L., M.W.A.D, L.T., B.J.M, S.J.F. and G.B. designed and planned the experimental work. H.M.H., D.C.B. and G.B. performed the research. H.M.H., D.C.B., P.M.L., M.W.A.D, L.T., B.J.M, S.J.F. and G.B. interpreted and analysed the data. H.M.H., D.C.B. and G.B. performed statistical analysis. H.M.H, P.M.L., G.B., B.J.M. and S.J.F. wrote the manuscript. All authors reviewed the manuscript.

## Acknowledgement

We would like to acknowledge Shelley Lampkin and Australian Phenomics Facility (APF) for the maintenance of the mouse colonies. We would also like to thank Dr Matthew Dixon and Prof Leann Tilley for assisting with ektacytometry and FRAP techniques and the associated analyses. We are also grateful for the assistance of the Microscopy Unit of the Macquarie University in the sample preparation and operation of the scanning electron microscope.

## Supporting Information Captions

**Table S1.**
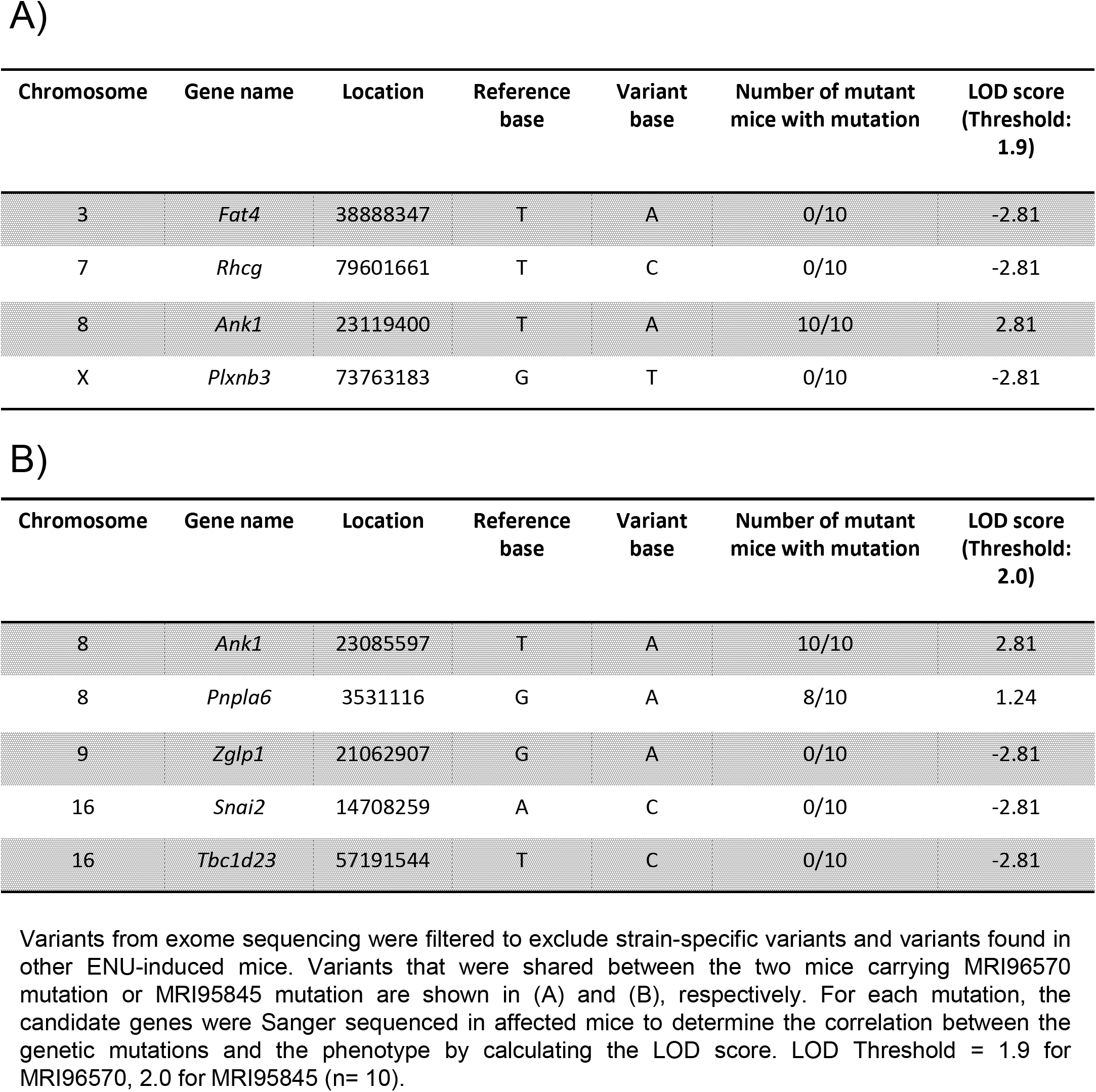
The candidate genes for MRI96570 and MRI95845 mutations. Variants from exome sequencing were filtered to exclude strain-specific variants and variants found in other ENU-induced mice. Variants that were shared between the two mice carrying MRI96570 mutation or MRI95845 mutation are shown in (A) and (B), respectively. For each mutation, the candidate genes were Sanger sequenced in affected mice to determine the correlation between the genetic mutations and the phenotype by calculating the LOD score. LOD Threshold = 1.9 for MRI96570, 2.0 for MRI95845 (n= 10).

| Chromosome | Gene name | Location | Reference base | Variant base | Number of mutant mice with mutation | LOD score (Threshold: 1.9) |
| --- | --- | --- | --- | --- | --- | --- |
| 3 | *Fat4* | 38888347 | T | A | 0/10 | -2.81 |
| 7 | *Rhcg* | 79601661 | T | C | 0/10 | -2.81 |
| 8 | *Ank1* | 23119400 | T | A | 10/10 | 2.81 |
| X | *Plxnb3* | 73763183 | G | T | 0/10 | -2.81 |

B)
| Chromosome | Gene name | Location | Reference base | Variant base | Number of mutant mice with mutation | LOD score (Threshold: 2.0) |
| --- | --- | --- | --- | --- | --- | --- |
| 8 | *Ank1* | 23085597 | T | A | 10/10 | 2.81 |
| 8 | *Pnpla6* | 3531116 | G | A | 8/10 | 1.24 |
| 9 | *Zglp1* | 21062907 | G | A | 0/10 | -2.81 |
| 16 | *Snai2* | 14708259 | A | C | 0/10 | -2.81 |
| 16 | *Tbc1d23* | 57191544 | T | C | 0/10 | -2.81 |

**Figure S1.**
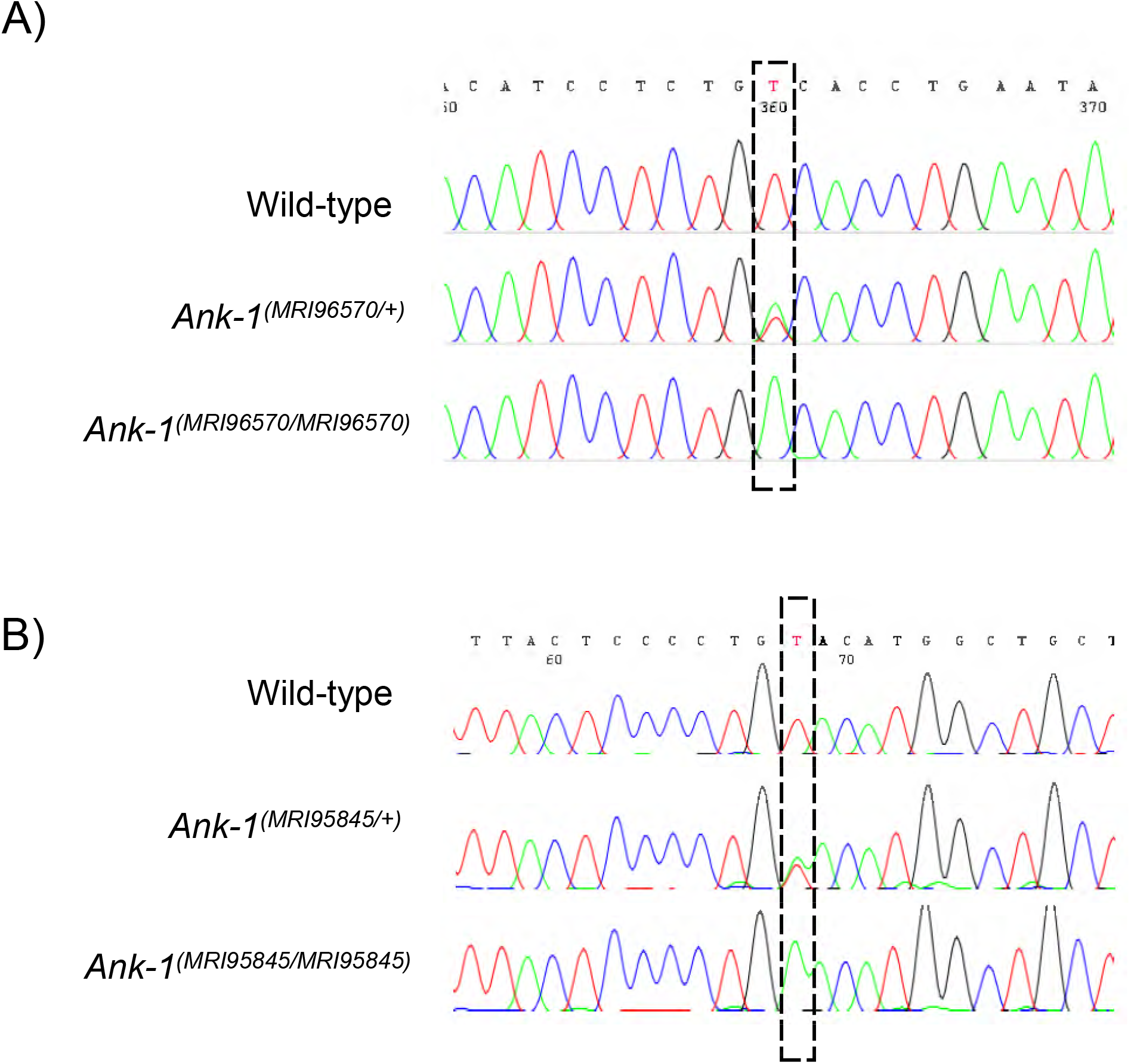
The location of *Ank-1^(MRI96570)^* and *Ank-1^(MRI95845)^* mutation. (A) Sanger sequencing of mice carrying *Ank-1^(MRI96570)^* revealed a T to A transversion in exon 34 of *Ank-1* gene, which is predicted to induce a premature stop codon. (B) Mice carrying *Ank-1^(MRI95845)^* mutation were found to have a T to A transversion in exon 5 of *Ank-1* gene, which is predicted to cause a missense mutation from tyrosine to asparagine at residue 149.

**Table S2.**
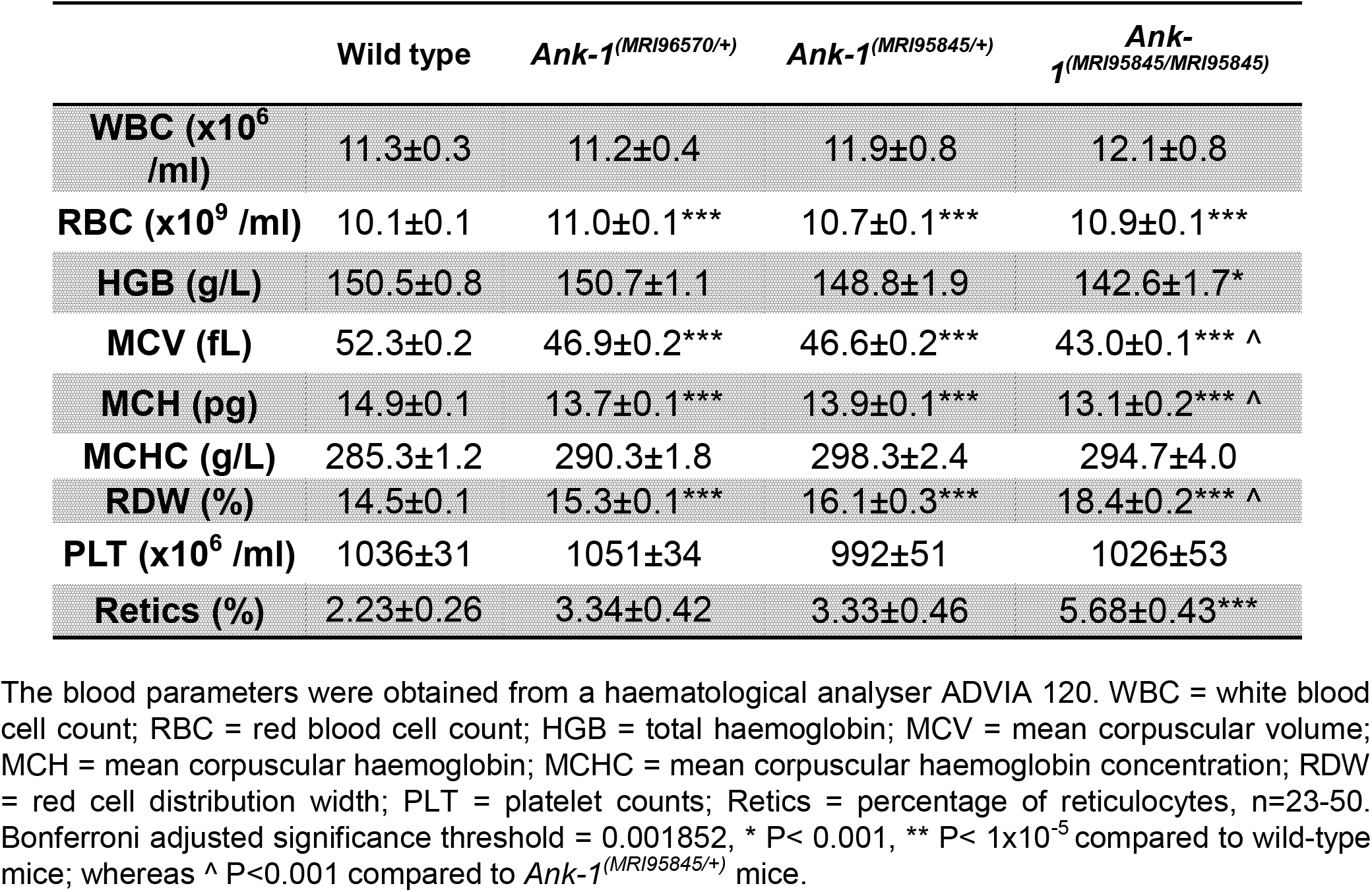
The complete blood count of *Ank-1^(MRI96570/+)^, Ank-1^(MRI95845/+)^* and *Ank-1^(MRI95845/MRI95845)^* mice. The blood parameters were obtained from a haematological analyser ADVIA 120. WBC = white blood cell count; RBC = red blood cell count; HGB = total haemoglobin; MCV = mean corpuscular volume; MCH = mean corpuscular haemoglobin; MCHC = mean corpuscular haemoglobin concentration; RDW = red cell distribution width; PLT = platelet counts; Reties = percentage of reticulocytes, n=23-50. Bonferroni adjusted significance threshold = 0.001852, * P< 0.001, ** P< 1×10^−5^ compared to wild-type mice; whereas ^^^ P<0.001 compared to *Ank-1^(MRI95845^*^/*+)*^ mice.

**Figure S2.**
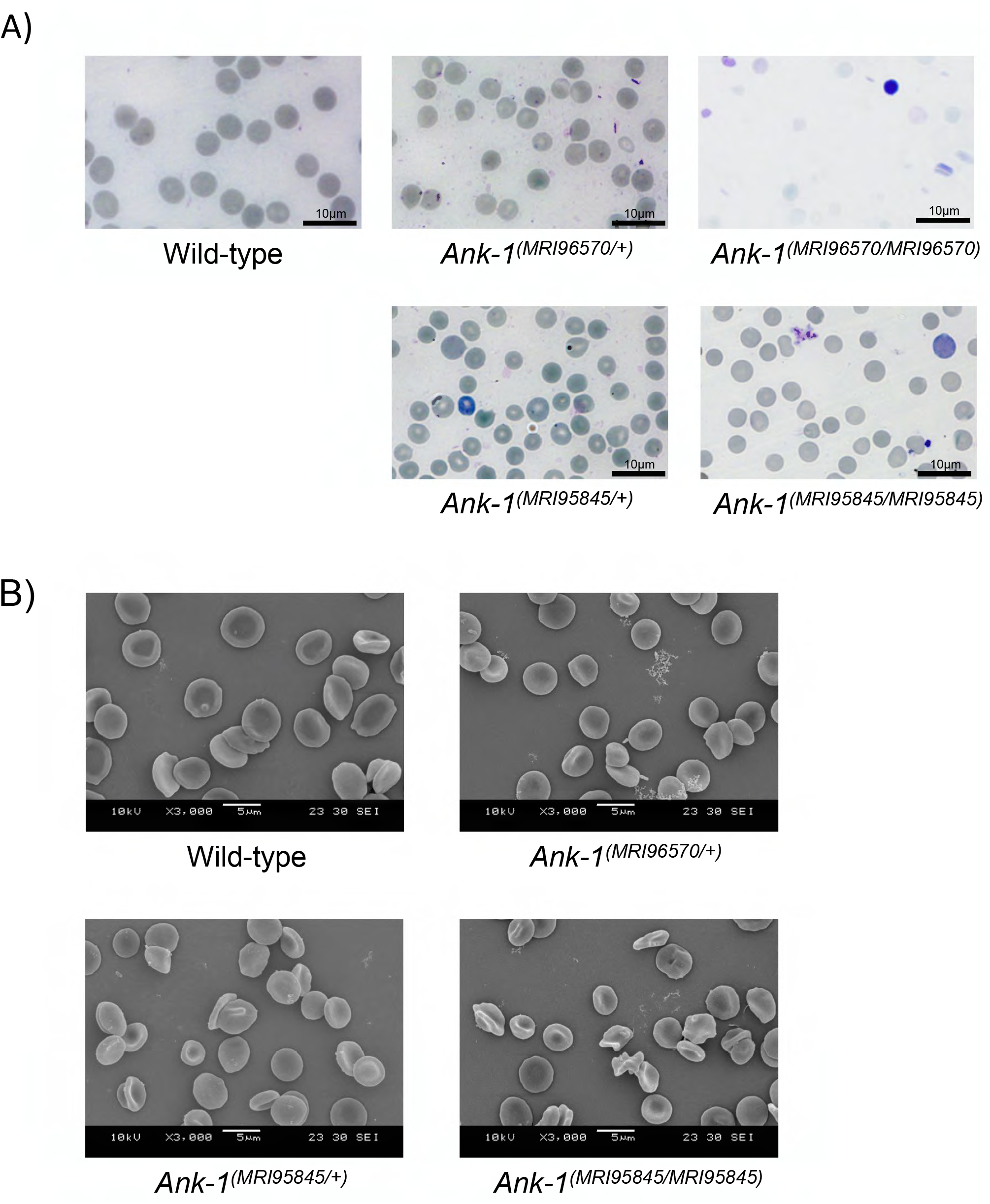
The RBC morphology of mice carrying *Ank-1^(MRI96570)^* or *Ank-1^(MRI95845)^* mutation. (A) Giemsa-stained blood smears as examined under light microscope at 1000x magnification. (B) Scanning electron microscopic images showing the RBC shape of *Ank-1^(MRI96570/+)^ Ank-1^(MRI95845)^* and *Ank-1^(MRI95845/MRI95845)^* mice.

**Figure S3.**
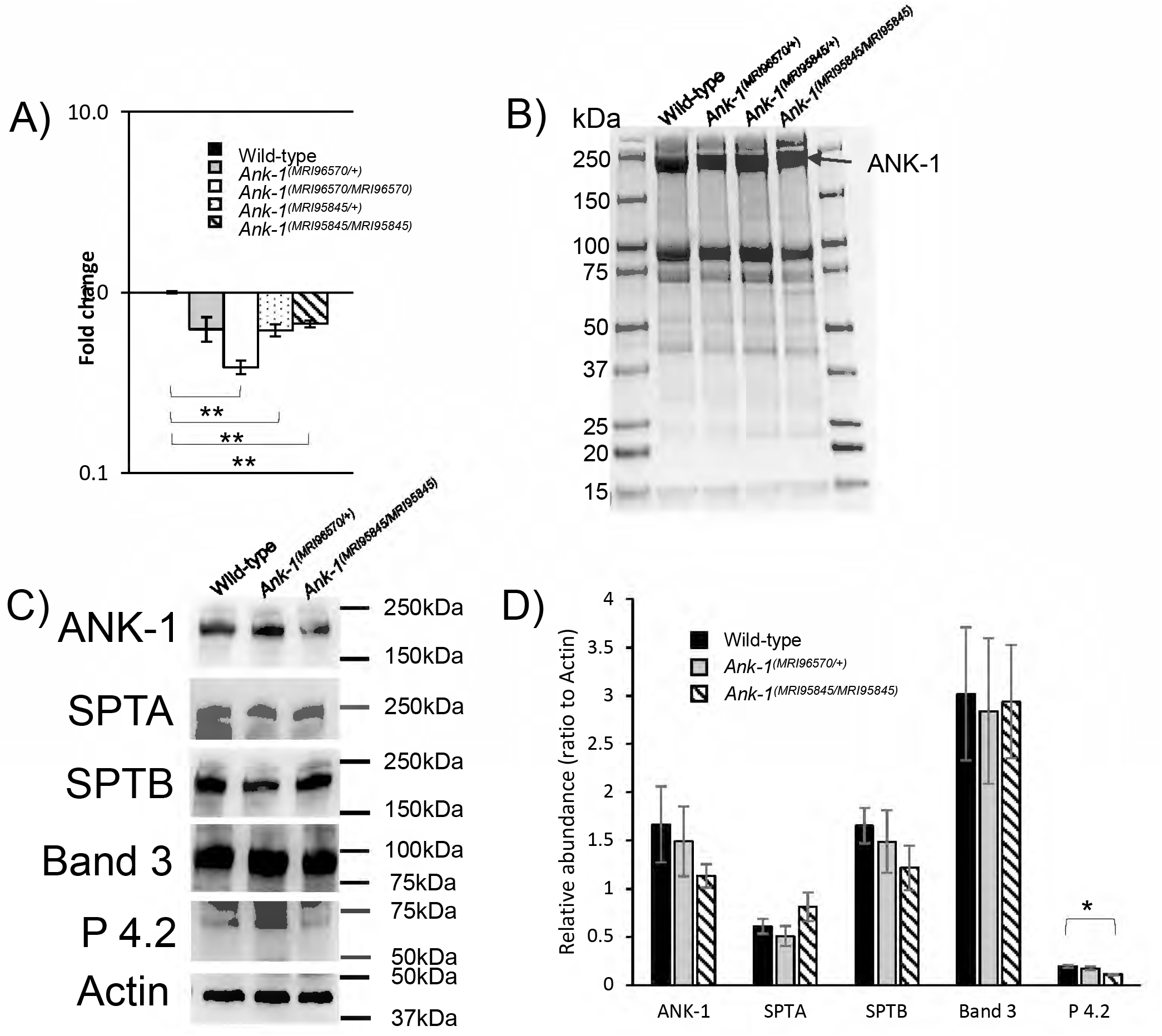
The expression of *Ank-1* and other RBC cytoskeletal proteins in mice carrying *Ank-1^(MRI96570)^* or *Ank-1^(MRI95845)^* mutation. (A) Quantitative PCR was carried out on E14 embryonic livers to examine *Ank-1* expression levels (n=3). The abundance of ANK-1 and other RBC cytoskeletal protein levels of *Ank-1^(MRI96570/+)^, Ank-1^(MRI95845/+)^* and *Ank-1^(MRI95845/MRI95845)^* mice as examined via (B) Coomassie staining and (C) Western blotting on membrane of mature RBCs. (D) The relative abundance of various cytoskeletal protein levels calculated from western blots (n=3). SPTA = α-spectrin, SPTB = β-spectrin, P 4.2 = Protein 4.2. * P<0.05, ** P<0.01. Error bars indicate SEM.

**Figure S4.**
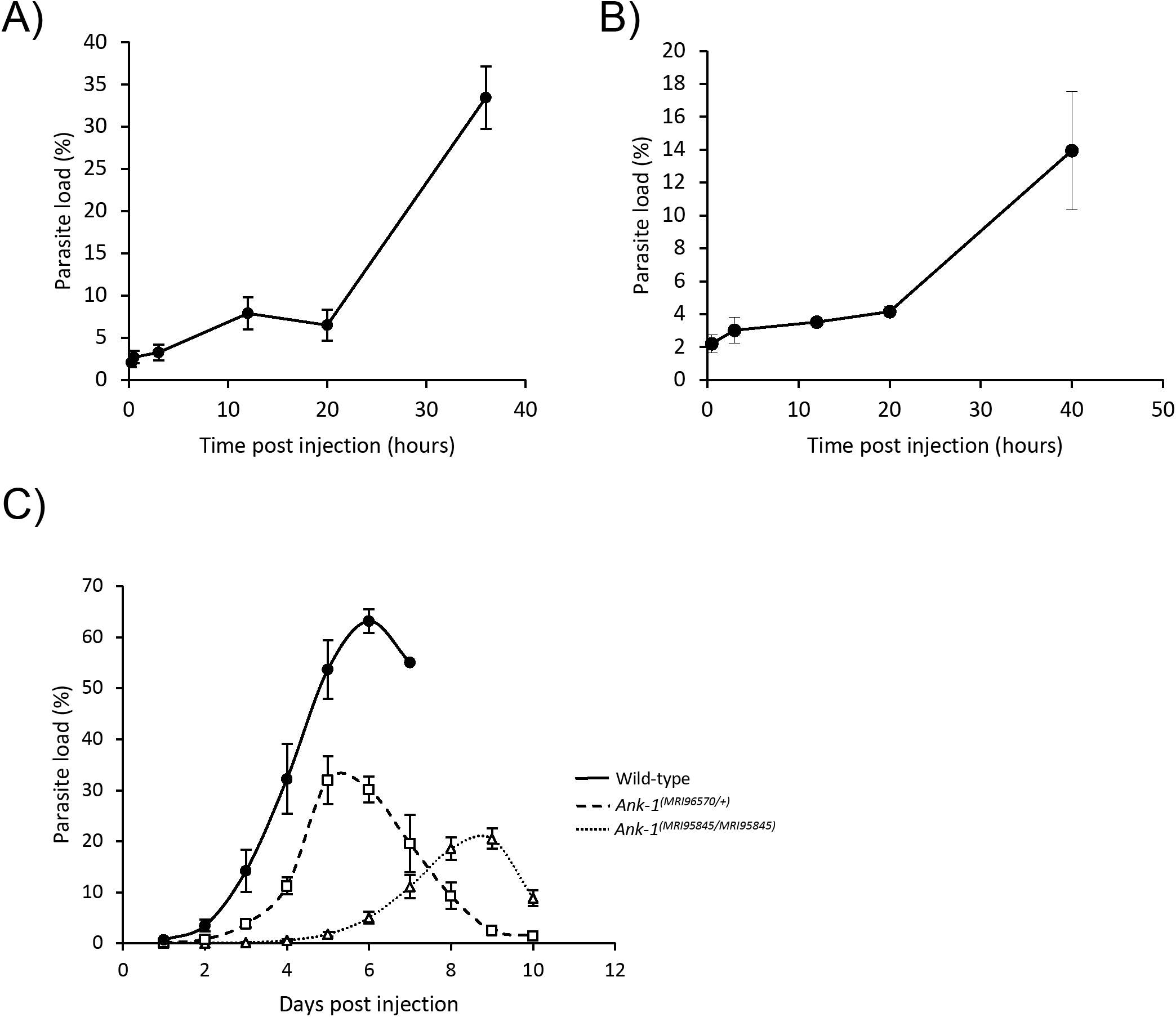
The parasitemia of the host mice during IVET assays and half-life assay. The parasitemia curve of the host mice during IVET assays, when comparing (A) wild-type with *Ank-1^(MRI95845/MRI95845)^* erythrocyteS; and (B) *Ank-1^(MRI96570/+)^* with *Ank-1^(MRI95845/MRI95845)^* erythrocytes (n=5-7). (C) The parasitemia curve of wild-type, *Ank-1^(MRI96570/+)^* and *Ank-1^(MRI95845/MRI95845)^* mice during half-life assay (n=6-7).

